# Heterogeneous complement and microglia activation mediates stress-induced synapse loss

**DOI:** 10.1101/2023.06.28.546889

**Authors:** Breeanne M. Soteros, Haven Tillmon, Mackenna Wollet, Julianne General, Hanna Chin, John Beichen Lee, Flavia R. Carreno, David A. Morilak, Jun Hee Kim, Gek Ming Sia

## Abstract

Spatially heterogeneous synapse loss is a characteristic of many psychiatric and neurological disorders, but the underlying mechanisms are unclear. Here, we show that spatially-restricted complement activation mediates stress-induced heterogeneous microglia activation and synapse loss localized to the upper layers of the mouse medial prefrontal cortex (mPFC). Single cell RNA sequencing also reveals a stress-associated microglia state marked by high expression of the apolipoprotein E gene (ApoE^high^) localized to the upper layers of the mPFC. Mice lacking complement component C3 are protected from stress-induced layer-specific synapse loss, and the ApoE^high^ microglia population is markedly reduced in the mPFC of these mice. Furthermore, C3 knockout mice are also resilient to stress-induced anhedonia and working memory behavioral deficits. Our findings suggest that region-specific complement and microglia activation can contribute to the disease-specific spatially restricted patterns of synapse loss and clinical symptoms found in many brain diseases.

## Introduction

Synapse loss is a central mechanism that plays an important role in the pathogenesis of many psychiatric and neurological diseases^1–6^. Neuroimaging and neuropathological studies have revealed disease-specific and reproducible spatial patterns of synapse loss in many brain disorders^7–10^. In some neuropsychiatric diseases, severe synapse loss can occur in remarkably small brain regions, while nearby regions are spared. For example, postmortem studies of schizophrenic patients show a consistent loss of dendritic spines in layer 3 of the dorsolateral prefrontal cortex (PFC)^11,12^, while spine density in deeper PFC layers^13^ and other cortical regions^14^ are unaffected. Studies of rodent models of depression also show that dendritic spines in the upper layers of the medial PFC (mPFC) are lost in response to stress, while spines in the deeper mPFC layers remain unaffected^15,16^. Despite the prevalence of region-specific synapse loss in brain diseases, the pathophysiological processes involved remain unclear.

Microglia are dynamic brain-resident macrophages that constantly survey their environment and respond to a wide variety of pathological insults. Single cell-RNA sequencing studies have revealed that under physiological conditions, microglia adopt diverse regional states dictated by their local environment^17–20^. A large number of pathological and spatially heterogeneous microglial transcriptomic states have also been described diseases such as Alzheimer’s disease^21–23^, multiple sclerosis^19,24,25^, and chronic pain^26^. These diverse microglia states are likely to perform key homeostatic functions in the healthy brain and play critical pathogenic roles in the diseased brain. While several signaling pathways have been shown to induce specific pathological microglial states^27–30^, whether these mechanisms act in a spatially-restricted manner remain unclear.

Stress is a major risk factor for psychiatric diseases, and extended periods of stress can directly cause affective disorders such as major depressive disorder (MDD)^31^, post-traumatic stress disorder (PTSD)^32,33^, and anxiety disorders^32,33^. Extended periods of stress also cause synapse loss in the PFC, an effect which is mediated by glucocorticoid stress hormones. Excessive secretion of cortisol is found in 50% of depressed patients^34^, and levels of cortisol are positively correlated to depressive behavior in non-human primates^35^. In humans, elevated salivary, hair and serum cortisol levels are all correlated with poorer working memory performance and lower brain volumes^36–39^, suggesting that prolonged exposure to cortisol can lead to changes in brain structure. Chronic stress is known to increase microglia activation in the PFC^40,41^, and can also induce complement activation in the brain^42–44^, suggesting that microglial phagocytosis of synapses may mediate stress-induced synapse loss. However, previous studies have assumed that stress-induced microglia activation occurs homogeneously throughout the PFC. Because the complement cascade is a secreted and extracellular signaling system, it was unclear whether this signaling pathway can result in spatially-restricted microglia activation and synapse loss in the brain.

Here, we provide evidence that chronic stress causes heterogeneous microglia activation in the mPFC resulting in layer-specific synapse loss and stress-associated behavioral deficits. We show that multiple chronic stress protocols cause localized complement activation specifically in the upper layers of the mouse mPFC. Complement activation in layer 2/3 leads to microglial phagocytosis of VGlut2 synapses and synapse loss specifically in those cortical layers, while synapses in deeper mPFC layers are spared. Single-cell RNA (scRNA-seq) of microglia from cortices of stressed mice also reveals an ApoE^high^ transcriptomic state upregulated in the stressed brain and localized to layer 2/3 of the mPFC. In complement C3 knockout (KO) mice, we found that chronic stress no longer induces the ApoE^high^ microglia state. C3 KO mice are also resilient to stress-induced microglia phagocytosis of synapses and synapse loss, and do not exhibit stress-induced anhedonia and working memory deficits. Our data shows that spatially restricted complement activation can drive heterogeneous activation of microglia to cause layer-specific synapse loss in the stressed mPFC.

## Results

### Multiple stress protocols cause layer-specific complement activation in the mPFC

The classical complement cascade is a major signaling pathway that mediates microglial phagocytosis of synapses^45–47^. Complement component C3, the main effector molecule in the complement cascade, is deposited onto synapses and recognized by microglial CR3 receptors for phagocytosis^45^. To determine whether chronic stress induces heterogeneous complement activation, we subjected C57BL/6J mice to the 21-day chronic corticosterone (CORT) administration protocol (Figure 1A), and assessed complement activation in the mPFC by staining for C3. We focused on the prelimbic cortex of the mPFC (Figure 1B), a region known to be especially sensitive to the effects of stress^48^. Surprisingly, we found that CORT treatment causes a layer-specific increase in C3 deposition in layer 2/3 of the mPFC but not layer 5/6 (Figures 1C-E). The effects of stress on neuroimmune processes can vary depending on the stressor used^49^, raising the question of whether layer-specific complement activation is a core feature of diverse stressors. In order to determine whether layer-specific complement activation also occurs in other stressors, we assessed C3 deposition in two other commonly used stress protocols, chronic restraint stress (CRS), and chronic unpredictable stress (CUS). Both of these protocols engage the fast autonomic nervous system (ANS) response, in addition to the slow hypothalamic-pituitary-adrenal (HPA) axis hormonal response that the CORT protocol mimics. Both of these protocols also provide different patterns of CORT responses. CRS is expected to cause an initial large increase in CORT levels which habituates over time, whereas CUS is designed to cause a non-habituating increase in CORT levels. Surprisingly, both CRS and CUS cause a layer-specific increase in C3 deposition similar to that found in chronic CORT administration (Figures 1F-K), despite differences in the physiological systems engaged and patterns of CORT induced by these protocols. Therefore, multiple chronic stress protocols lead to layer-specific complement activation in the mPFC, suggesting that layer-specific complement activation is a core feature of chronic stress.

**Figure 1.**
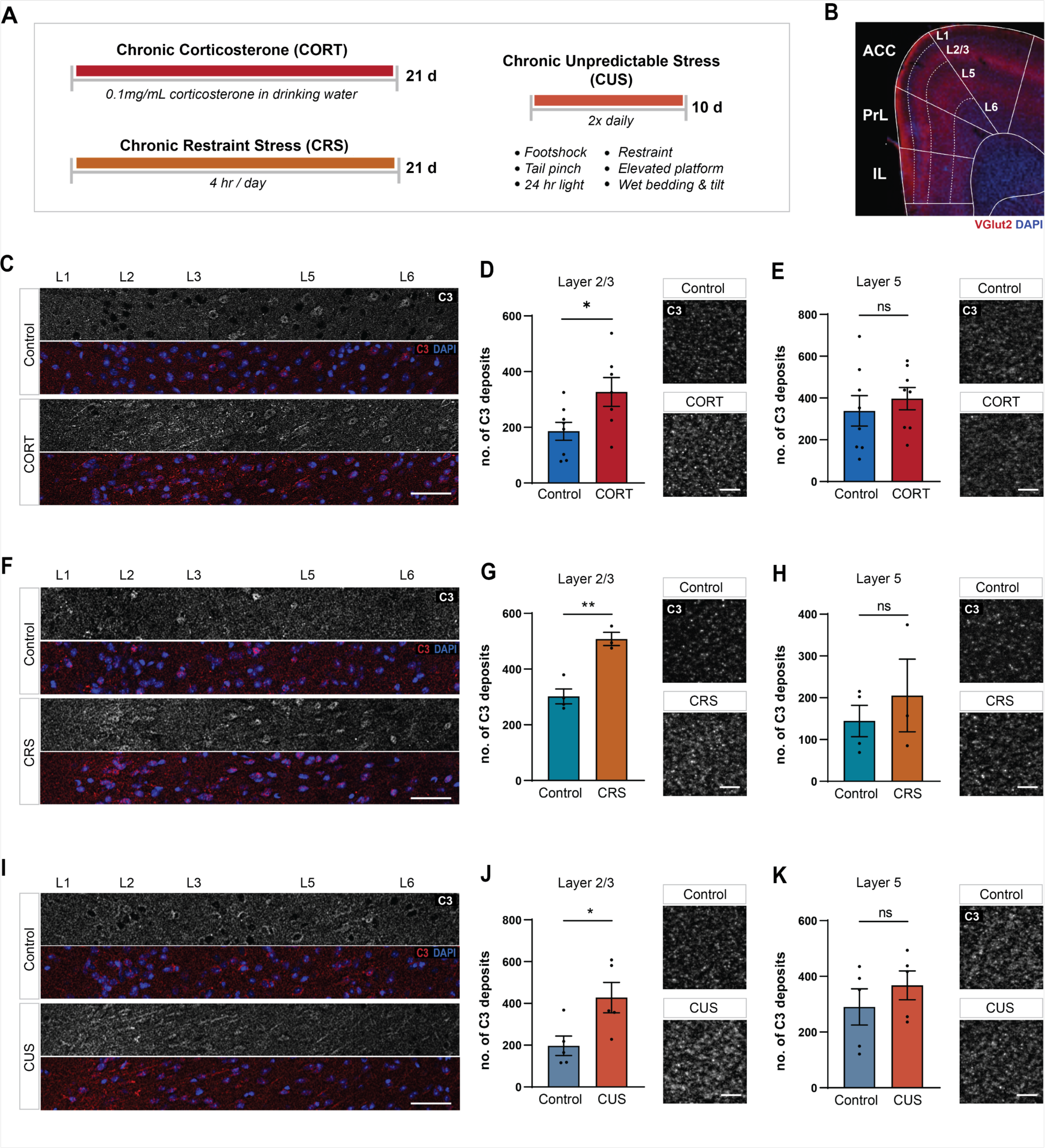
Multiple stress protocols cause layer-specific complement activation in the mPFC. (A) Schematic depicting timeline for chronic corticosterone (CORT) stress, chronic restraint stress (CRS), and chronic unpredictable stress (CUS) protocols. (B) Low-magnification image depicting layers of the medial prefrontal cortex. Stained for VGlut2 (red) and DAPI (blue). (C) Low magnification image showing laminar C3 deposition in mPFC in control/CORT mice. Top panel shows C3 channel (white), bottom panel shows merged C3 (red) and DAPI (blue) channels. Scale bar 50 µm. (D) Quantitation (left) and representative image (right) of C3 deposition in layer 2/3 mPFC after control/CORT. Analyzed by two-tailed unpaired t-test, t(13)=2.394, *p=0.0325, n=7-8 from 3-4 mice. All data shown as mean ± s.e.m.. Scale bar 5 µm. (E) Quantitation (left) and representative image (right) of C3 deposition in layer 5 mPFC after control/CORT. Analyzed by two-tailed unpaired t-test, t t(14)=0.6462, p=0.5286, n=8 from 4 mice. All data shown as mean ± s.e.m.. Scale bar 5 µm. (F) Low magnification image showing laminar C3 deposition in mPFC in control/CRS mice. Top panel shows C3 channel (white), bottom panel shows merged C3 (red) and DAPI (blue) channels. Scale bar 50 µm. (G) Quantitation (left) and representative image (right) of C3 deposition in layer 2/3 mPFC after control/CRS. Analyzed by two-tailed unpaired t-test, t(5)=5.491, **p=0.0027, n=3-4 mice. All data shown as mean ± s.e.m.. Scale bar 5 µm. (H) Quantitation (left) and representative image (right) of C3 deposition in layer 5 mPFC after control/CRS. Analyzed by two-tailed unpaired t-test, t(5)=0.7149, p=0.5066, n=3-4 mice. All data shown as mean ± s.e.m.. Scale bar 5 µm. (I) Low magnification image showing laminar C3 deposition in mPFC in control/CUS mice. Top panel shows C3 channel (white), bottom panel shows merged C3 (red) and DAPI (blue) channels. Scale bar 50 µm. (J) Quantitation (left) and representative image (right) of C3 deposition in layer 2/3 mPFC after control/CUS. Analyzed by two-tailed unpaired t-test, t(8)=2.675, *p=0.0281, n=5 from 3 mice. All data shown as mean ± s.e.m.. Scale bar 5 µm. (K) Quantitation (left) and representative image (right) of C3 deposition in layer 5 mPFC after control/CUS. Analyzed by two-tailed unpaired t-test, t(8)=0.9416, p=0.3739, n=5 from 3 mice. All data shown as mean ± s.e.m.. Scale bar 5 µm

### Stress causes layer-specific microglial synapse phagocytosis and synapse loss

Complement activation is known to lead to microglial activation and phagocytosis of synapses^45^. To further characterize microglia in the context of chronic stress, we chose to perform all further experiments with the CORT stress protocol, because CORT is the principal murine stress hormone, and CORT levels are increased by all rodent stress protocols^50–52^. Furthermore, direct CORT administration causes synapse losses and behavioral deficits in rodents^53–57^, and glucocorticoid receptor blockade in rodents block stress-induced microglia activation, neuronal remodeling, and behavioral deficits^57^, indicating that CORT is necessary and sufficient for the neurobiological effects of chronic stress. To assess microglial activation, we stained brain slices for the microglial cytoplasmic marker Iba1 and the lysosomal marker CD68 (Figure 2A), reconstructed high resolution 3D images of individual microglia, and quantitated the percentage of Iba1 volume occupied by CD68 lysosomes. We found that the CD68 content of microglia is increased by chronic CORT administration only in layer 2/3 (Figure 2B), while microglia in layer 5 are not affected (Figure 2C). Previous studies have shown that chronic CORT administration causes synapse loss in the mPFC^53–55^. Furthermore, live imaging studies indicate that stress-induced synapse loss in the mPFC is due to an increase in the rate of synapse elimination^54,58,59^, as opposed to a decrease in the rate of synapse formation, suggesting that microglial synapse phagocytosis may be involved. To assess microglial synapse phagocytosis in the mPFC of chronically stressed mice, we stained their brain slices for Iba1, CD68, and the presynaptic marker VGlut2. We chose VGlut2 because studies from our lab and others have shown that microglia preferentially phagocytose this population of synapses^45,46,60^. We quantitated the 3D colocalization of the three markers Iba1/CD68/VGlut2 as a measure of microglial synapse phagocytosis (Figure 2D), and found that chronic CORT administration strongly increases microglial synapse phagocytosis in layer 2/3 of the mPFC, but not in layer 5/6 (Figures 2E-F).

**Figure 2.**
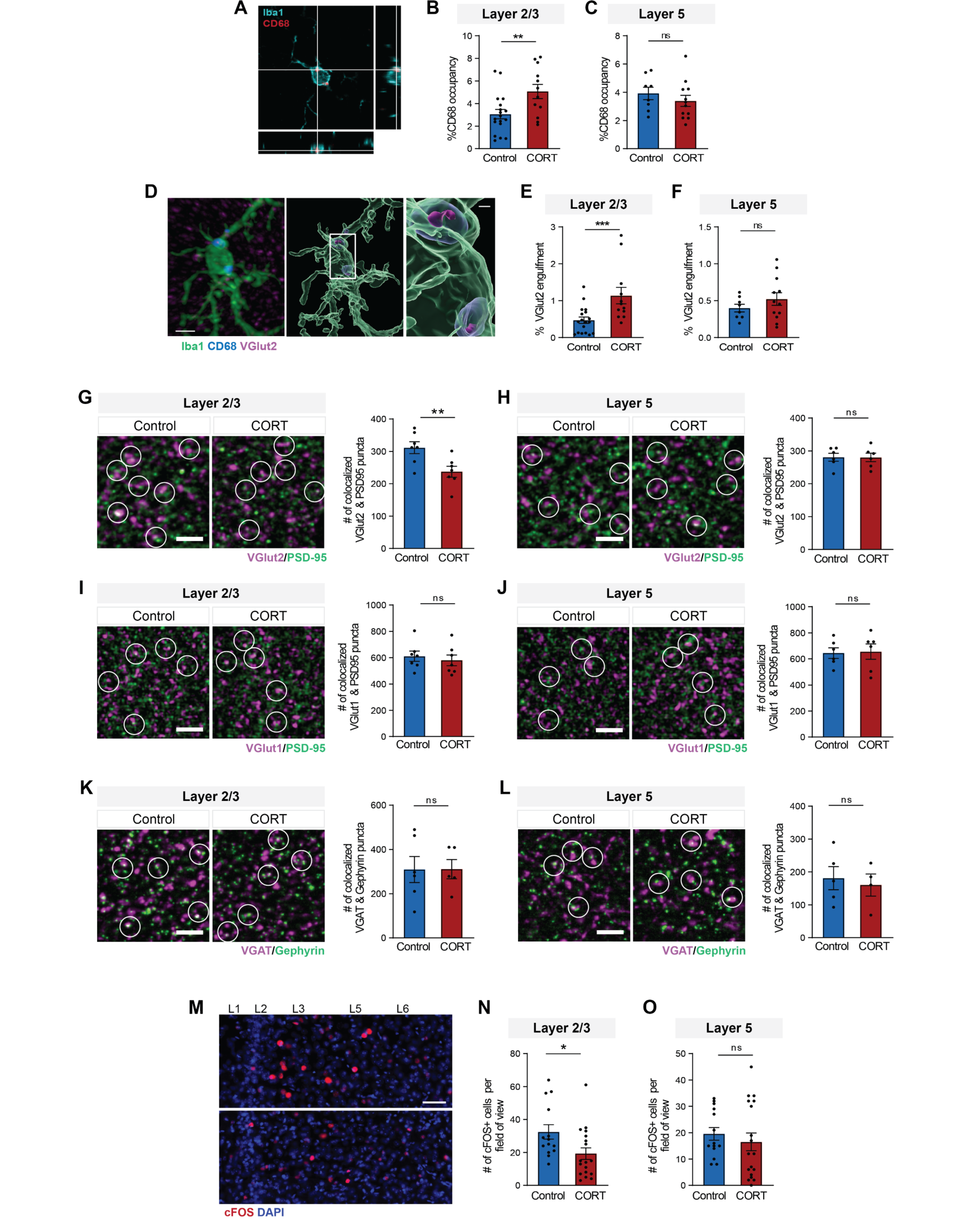
Chronic stress causes layer-specific microglial activation and synapse loss in mPFC. (A) Orthogonal view showing localization of lysosomal marker CD68 (red) within Iba1-positive microglia (blue). (B) CD68 occupancy of microglia in mPFC PrL layer 2/3. Analyzed by two-tailed unpaired t-test, t(28)=2.790, **p=0.0094, n=12-18 cells from 4 mice. (C) CD68 occupancy of microglia in mPFC PrL layer 5. Analyzed by two-tailed unpaired t-test, t(18)=0.8727, p=0.3943, n=8-12 cells from 3-4 mice. All data shown as means ± s.e.m.. (D) Microglial engulfment assay. Left: Representative IHC staining of Iba1-positive microglia (green) with CD68-positive lysosomes (blue) and VGlut2 (magenta), scale bar 5 µm. Right: Corresponding 3D reconstruction, with inset highlighting engulfed VGlut2 within microglial lysosomes. Inset scale bar 1 µm. (E) VGlut2 engulfment by microglia in mPFC PrL layer 2/3. Analyzed by two-way ANOVA, followed by Sidak’s post-hoc test comparing means within genotype, F(1,44)=6.067, p=0.0178, n=12-18 cells from 4 mice, ***p=0.0006. Data excerpted from wildtype groups in Figure 6D. (F) VGlut2 engulfment by microglia in mPFC PrL layer 5. Analyzed by two-way ANOVA, F(1,31)=0.6694, p=0.4094, n=8-12 cells from 3-4 mice. All data shown as means ± s.e.m. Data excerpted from wildtype groups in Figure 6F. (G) VGlut2/PSD95 synapse density in layer 2/3 mPFC. Representative images (left) and quantitation (right) of colocalized VGlut2 (magenta) and PSD95 (green) puncta. Circles indicate colocalized pre- and post-synaptic puncta. Scale bar 5 µm. All data shown as mean ± s.e.m. Analyzed by two-way ANOVA, followed by Sidak’s post-hoc test comparing means within genotype, F(1,24)=10.67, p=0.0033 (genotype x stress interaction), n=7 from 3 mice, **p=0.0043. Data excerpted from wildtype groups in Figure 6C. (H) VGlut2/PSD95 synapse density in layer 5 mPFC. Analyzed by two-way ANOVA, F(1,21)=0.07829, p=0.7824, n=6-7 from 3 mice. Data excerpted from wildtype groups in Figure 6E. (I) VGlut1/PSD95 synapse density in layer 2/3 mPFC. Representative images (left) and quantitation (right) of colocalized VGlut1 (magenta) and PSD95 (green) puncta. Circles indicate colocalized pre- and post-synaptic puncta. Scale bar 5 µm. All data shown as mean ± s.e.m. Analyzed by two-tailed unpaired t-test, t(12)=0.5547, p=0.5893, n=7 from 3 mice. (J) VGlut1/PSD95 synapse density in layer 5 mPFC. Analyzed by two-tailed unpaired t-test, t(10)=0.1575, p=0.8780, n=6 from 3 mice. (K) VGAT/gephyrin synapse density in layer 2/3 mPFC. Representative images (left) and quantitation (right) of colocalized VGAT (magenta) and gephyrin (green) puncta. Circles indicate colocalized pre- and post-synaptic puncta. Scale bar 5 µm. All data shown as mean ± s.e.m. Analyzed by two-tailed unpaired t-test, t(9)=0.01659, p=0.9871, n=5-6 from 3 mice. (L) VGAT/gephyrin synapse density in layer 5 mPFC. Analyzed by two-tailed unpaired t-test, t(7)=0.4156, p=0.6901, n=4-5 from 2-3 mice. (M) Representative low-magnification image of c-fos (red) and DAPI (blue) staining in mPFC after TOR working memory task. Scale bar 50 µm. (N) Quantitation of c-fos-positive neurons in layer 2/3 mPFC after TOR task. Analyzed by two- tailed unpaired t-test, t(30)=2.371, *p=0.0244, n=14-18 from 3-4 mice. Data shown as mean ± s.e.m. (O) Quantitation of c-fos-positive neurons in layer 5 mPFC after TOR task. Analyzed by two-tailed unpaired t-test, t(30)=0.6996, p=0.4896, n=14-18 from 3-4 mice. Data shown as mean ± s.e.m.

To determine if the increase in microglial synapse phagocytosis leads to a decrease in synapse density in the stressed mPFC, we stained brain slices for three types of synapses: 1) thalamocortical synapses marked by colocalized VGlut2 presynaptic and PSD95 postsynaptic markers, 2) corticocortical synapses marked by colocalized VGlut1 presynaptic and PSD95 postsynaptic markers, and 3) inhibitory synapses marked by colocalized VGAT presynaptic and gephyrin postsynaptic markers. We found that chronic CORT administration results in a layer-specific reduction of VGlut2/PSD95 synapses in layer 2/3, but not layer 5 (Figures 2G-H), while the densities of VGlut1/PSD95 and VGAT/gephyrin synapses are unchanged in all cell layers (Figures 2I-L). These findings suggest that microglial synapse phagocytosis induced by chronic CORT administration specifically targets thalamocortical synapses in layer 2/3 of the mPFC, while sparing corticocortical and inhibitory synapses.

Activity of the thalamic projections to the PFC is known to be required for working memory performance, where it acts to sustain the delay-period activity of PFC neurons which are the neuronal substrates of working memory maintenance^61–63^. Chronic stress is also known to impair thalamic inputs to the mPFC and working memory performance in mice^56,57,64^. To determine whether stress-induced loss of synapses in the mPFC results in a layer-specific reduction of functional activation of neurons during working memory maintenance, we utilized the temporal object recognition (TOR) working memory task, which has previously been shown to be impaired by chronic stress in mice^56,57^. We assessed the number of c-fos positive neurons in the mPFC shortly after the completion of the TOR task, and found a significant reduction in layer 2/3 of the mPFC from CORT mice, but not in layer 5/6 (Figures 2M-O). Together, these data indicate that chronic CORT causes heterogeneous microglia activation in the mPFC leading to layer-specific synapse loss.

### Stress induces an ApoE^high^ microglia state enriched in endocytotic genes

To characterize the heterogeneous states of microglia in the stressed brain, we performed scRNA-seq on microglia isolated from the brains of CORT and control mice. To isolate single cell microglia from the gray matter of mouse cortex, we used a cold mechanical dissociation protocol that was specifically optimized for the isolation of microglia with minimal *ex vivo* transcriptional alterations (Figure 3A)^19,65^. After dissociation, microglia were sorted from the single cell suspension with fluorescence activated cell sorting (FACS) based on expression of Cd45, Cd11b, and Cx3cr1. (Supplementary Figure 1). The resulting cells were processed for single cell RNA sequencing, and sequenced to a depth of more than 50,000 reads/cell. We processed 2 replicates each for the control and CORT conditions, with 4 mice per replicate. After quality control, we retained a total of 35,292 cells from all replicates for downstream analysis.

**Figure 3.**
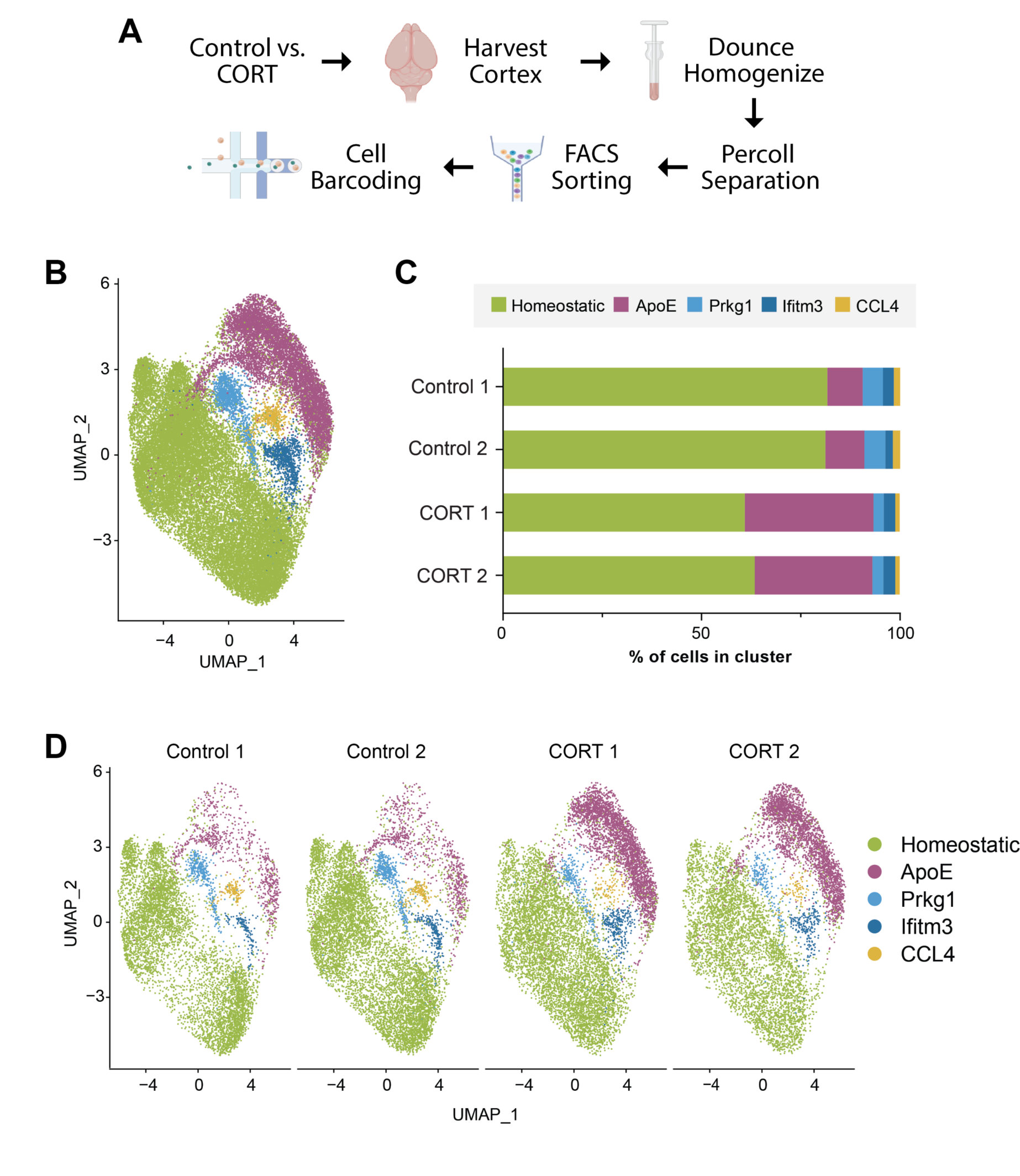
Chronic CORT administration induces an ApoE^high^ microglia state. (A) Schematic of microglia single cell isolation, FACS, and single cell RNA sequencing process. (B) UMAP plot of 35,292 microglia isolated from the cerebral cortex of control and CORT mice, 2 replicates per condition, n = 4 mice per replicate. In total, 5 clusters of microglia were identified. The largest cluster was annotated the homeostatic cluster, while other clusters were annotated by the most differentially expressed gene compared to other clusters. (C) Proportion of cells in each cluster for all replicates of both control and CORT conditions. (D) UMAP plots of cells from each replicate.

We then performed unsupervised clustering analysis on the single scRNA-seq dataset, revealing 5 clusters across all conditions and replicates (Figure 3B). We analyzed each cluster for differentially expressed genes (DEGs), and annotated each cluster with its most highly expressed DEG, with the exception of the largest cluster, which we termed the homeostatic cluster (Figures 3C, D). In accordance with previous reports, adult microglia are largely homogeneous and are mostly in the homeostatic state (Figures 3B, C)^18,19,65^. Chronic CORT administration dramatically increases the number of microglia in the ApoE^high^ state in both CORT biological replicates (Figures 3C, D), while decreasing the number of homeostatic microglia. Compared to the homeostatic state, the ApoE^high^ microglia state exhibits greatly increased expression of the ApoE gene, with a nearly 32-fold increase over the homeostatic state, as well as increases in the expression of several Ms4a genes (Figure 4D). Intriguingly, both ApoE and Ms4a genes are associated with AD risk^66–68^, suggesting a microglia-dependent mechanism for acceleration of AD progression by stress. In addition to the homeostatic and ApoE^high^ cell clusters, we also identified several smaller cell clusters which are largely unchanged by chronic CORT administration (Figures 3C, D), many of which have already been identified in previous microglia scRNA-seq studies^65,69,70^, suggesting that they are microglia states present in basal conditions. These clusters include an interferon-responsive state marked by expression of Ifitm3 and other interferon-responsive genes^65,70^, and a chemokine-expressing state marked by Ccl4 and other chemokine expression (Figures 3B, C)^65,69^. We also identified a microglia state marked by expression of the protein kinase Prkg1, which to our knowledge has not been previously described (Figures 3B, C). All cell clusters express canonical microglial genes such as C1qa, Cx3cr1, P2ry12, and Tmem119 (Figure 4A), indicating that they are *bona fide* microglia states.

**Figure 4.**
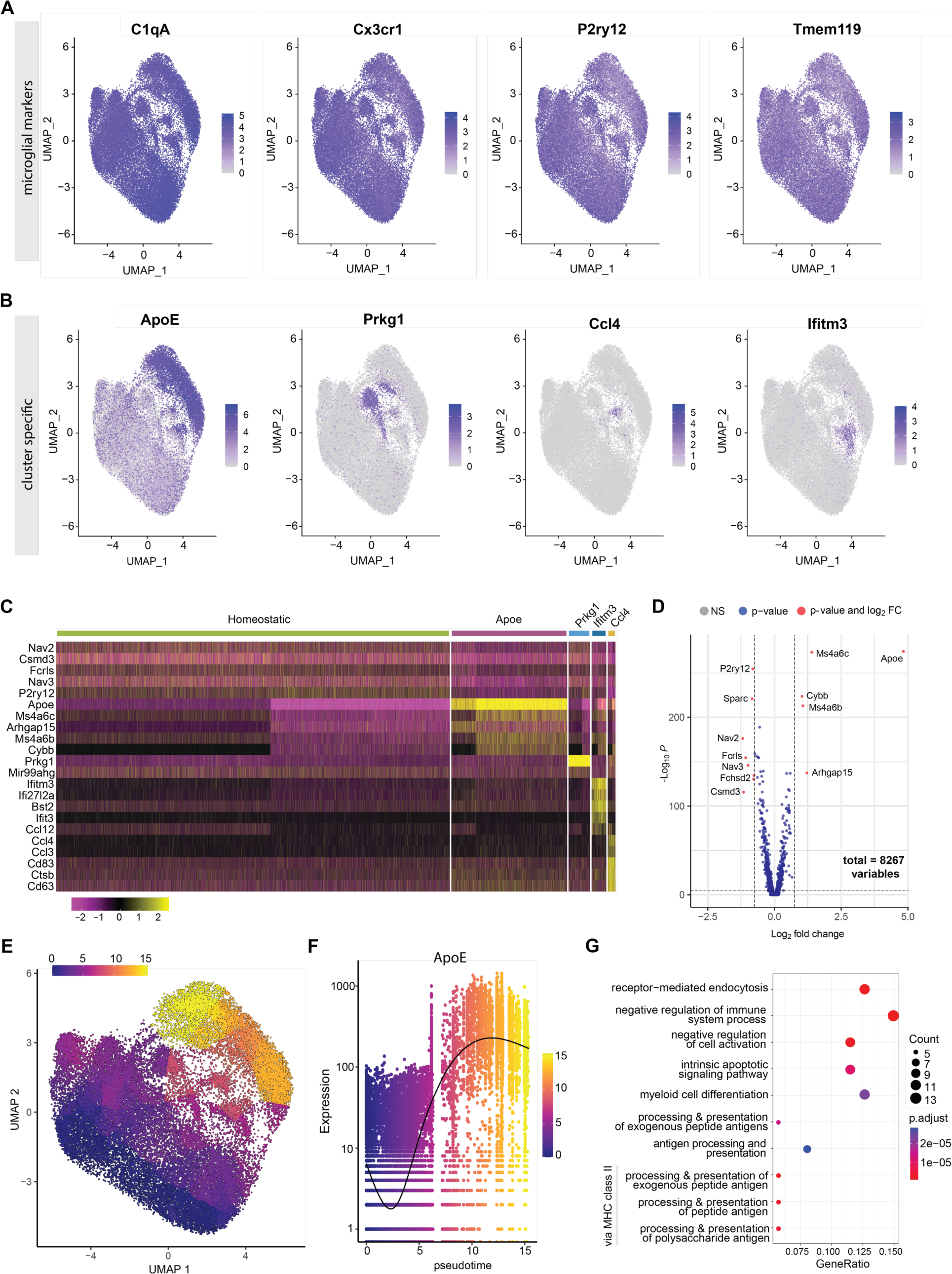
Endocytosis-associated genes are upregulated in the ApoE^high^ microglia state. (A) UMAP plots showing expression of canonical microglial genes. (B) UMAP plots showing expression of marker genes for the Apoe, Prkg1, Ccl4, and Ifitm3 clusters. (C) Heat map showing the top genes that are upregulated in each cluster relative to all other clusters. (D) Volcano plot of differentially expressed genes (DEGs) of the Apoe^hi^ cluster versus the homeostatic cluster. Genes with log_2_-fold change > 0.75 are labeled. (E) Pseudotime distribution of microglia in UMAP plot, with the homeostatic cluster designated as the root of the trajectory. (F) Expression level of the ApoE gene in pseudotime. (G) Dot plot showing top 10 biological function GO terms identified by gene set enrichment analysis (GSEA) of DEGs from pseudotime analysis. GO terms are arranged in order of increasing adjusted p value.

Chronic CORT administration increases the number of microglia in the ApoE^high^ state while decreasing the number of homeostatic microglia (Figures 3B, C), suggesting that CORT induces microglia to convert from the homeostatic state to the ApoE^high^ state. To identify the genes involved in this transition, we performed pseudotime trajectory analysis on the dataset. This analysis corroborated that the homeostatic population transitioned to the ApoE^high^ population along a single trajectory (Figure 4E), and that 93 genes were differentially expressed along this trajectory, including the ApoE gene (Figure 4F). We then performed gene set enrichment analysis (GSEA) on the 93 genes, which showed that genes associated with receptor mediated endocytosis were significantly enriched among these genes (Figure 4G). In addition, genes associated with negative regulation of T cell activation and antigen presentation were also differentially expressed in microglia (Figure 4G), consistent with the known immunosuppressive function of the glucocorticoids. The enrichment of genes involved in receptor mediated endocytosis suggests that ApoE^high^ microglia are primed for phagocytosis by stress.

### Complement activation drives formation of ApoE^high^ microglia state

Next, we wanted to determine if layer-specific complement activation resulted in layer-specific induction of ApoE^high^ microglia in the mPFC. We mapped ApoE-expressing microglia in the mPFC, using RNAscope *in situ* hybridization (ISH) to probe for both Cx3cr1 mRNA transcripts as a microglia marker, and ApoE transcripts as the readout. We took a semi-quantitative approach that was suggested by the manufacturer, categorizing Cx3cr1-positive microglia into bins based on their ApoE mRNA content, ranging from no expression (bin 0) to high expression (bin 4) (Figure 5A). We then characterized the bin distribution for ApoE expression in wildtype and C3^-/-^ mice treated with control or CORT (Figures 5B-E). Notably, in wildtype mice treated with CORT, over 25% of all microglia in layer 2/3 are bin 4 ApoE^high^ microglia, whereas less than 10% of microglia in control mice are bin 4 ApoE^high^ (Figures 5B, D). This dramatic increase in ApoE^high^ microglia population is not seen in layer 5 (Figures 5C, E). In addition, C3 deletion abolishes the layer-specific increase in ApoE^high^ microglia (Figures 5B-E). These data suggest that complement activation drives the formation of the ApoE^high^ microglia state.

**Figure 5.**
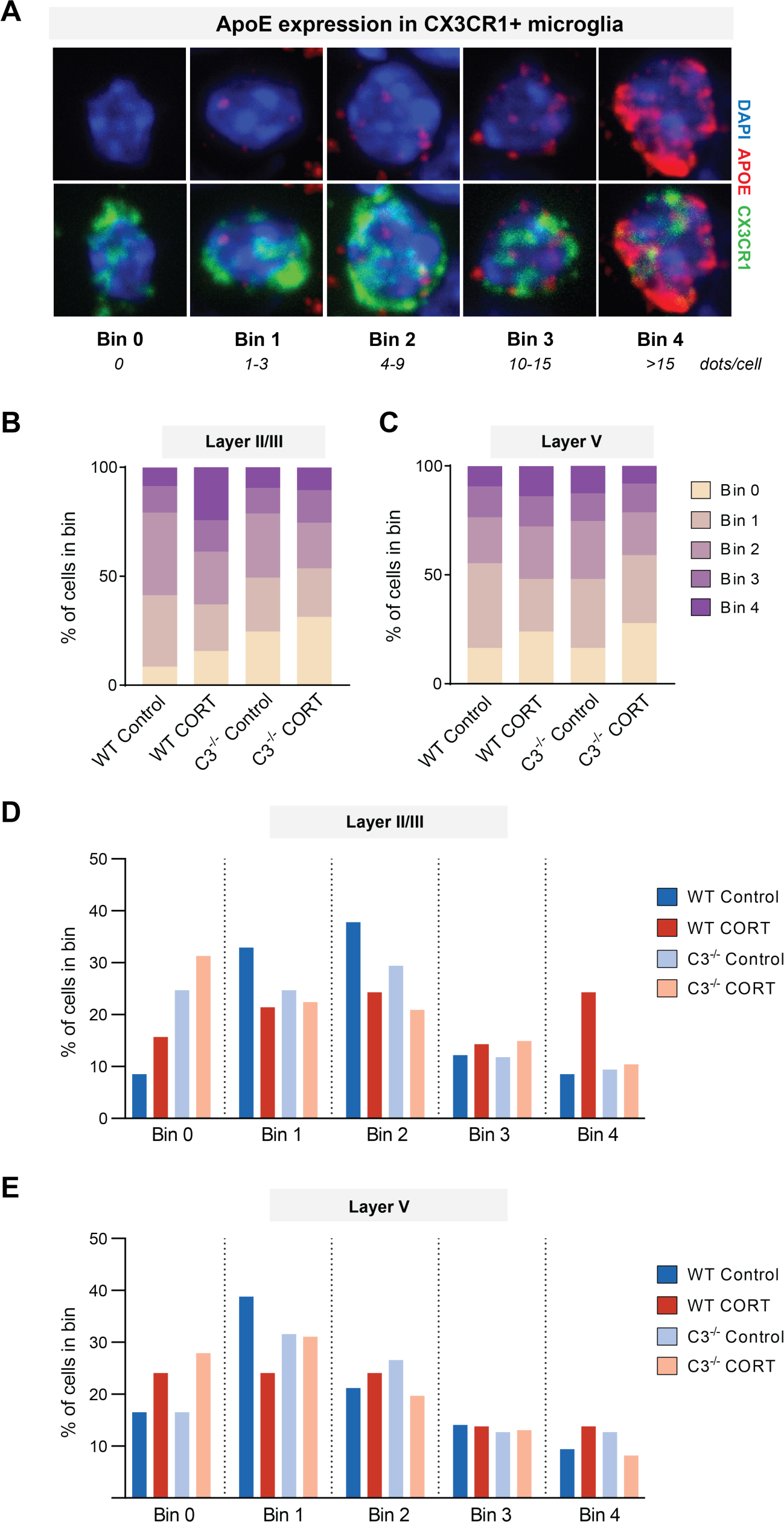
Complement activation drives formation of ApoE^high^ microglia state. (A) Semi-quantitative scoring of RNAscope *in situ* hybridization for ApoE mRNA (red) expression in Cx3cr1-positive microglia (green). Representative images of microglia categorized into bins (0-4) based on the number of ApoE puncta/clusters per nuclei. (B) Stacked bar graph showing ApoE bin distribution of microglia in layer 2/3 mPFC in WT and C3^-/-^ control and CORT mice, displayed as % of total cells in bin; 67-85 total cells per group from 2-3 mice per group. (C) Stacked bar graph showing ApoE bin distribution of microglia in layer 5/6 mPFC in WT and C3^-/-^ control and CORT mice, displayed as % of total cells in bin; 58-85 total cells per group from 2-3 mice per group. (D) Bar graph showing percentage of microglia within each ApoE bin category for each genotype in layer 2/3. (E) Bar graph showing percentage of microglia within each ApoE bin category for each genotype in layer 5/6.

### Complement activation drives stress-induced synapse loss and behavioral deficits

To determine if C3 deletion leads to a lack of stress-induced microglial and synaptic responses, we next assessed VGlut2/PSD95 synapse density and microglial phagocytosis of VGlut2 synapses in the mPFC of control and CORT-treated C3^-/-^ mice (Figures 6A, B). We found that mice lacking C3 were protected against CORT-induced microglial phagocytosis of synapses and synapse loss in layer 2/3 (Figures 6C-F). These data indicate that stress-induced layer-specific changes in microglial synapse phagocytosis and synapse density are abolished in the C3^-/-^ mice.

**Figure 6.**
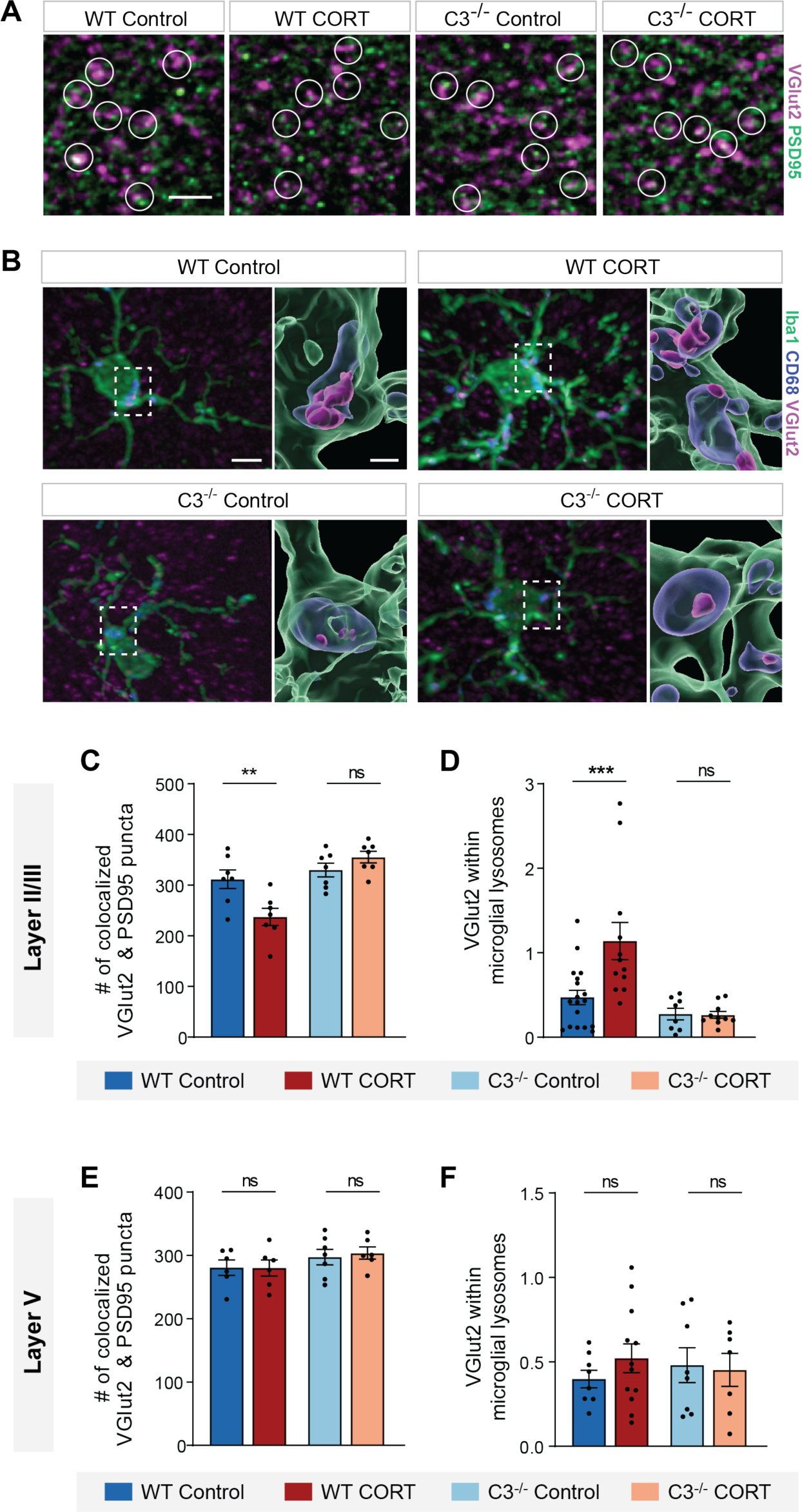
C3^-/-^ mice are protected from stress-induced synapse loss. (A) Representative images of VGlut2 (magenta) and PSD95 (green) synapses colocalization in layer 2/3 mPFC from wildtype and C3^-/-^ control/CORT mice. Circles indicate colocalized pre- and postsynaptic puncta. Scale bar 5 µm. (B) Representative images of microglial VGlut2 synapse engulfment in layer 2/3 mPFC from wildtype and C3^-/-^ control/CORT mice. Representative IHC image (left) with corresponding 3D reconstruction (inset, right) showing localization of engulfed VGlut2 (magenta) in CD68-positive lysosomes (blue) in Iba1-positive microglia (green). Scale bar 5 µm for IHC image, scale bar 1 µm for inset. (C) Quantitation of colocalized VGlut2 & PSD95 synaptic puncta in layer 2/3 mPFC of wildtype and C3^-/-^ control/CORT mice. Data analyzed by two-way ANOVA, followed by Sidak’s post-hoc test comparing means within genotype. F(1,24)=10.67, p=0.0033 (genotype x stress interaction), n=7 from 3 mice, **p=0.0043. Data plotted as mean ± s.e.m. (D) Quantitation of microglial VGlut2 engulfment in layer 2/3 mPFC of wildtype and C3^-/-^ mice. Data analyzed by two-way ANOVA, followed by Sidak’s post-hoc test comparing means within genotype, F(1,44)=6.067, p=0.0178 (genotype x stress interaction), n=12-18 cells from 4 mice, ***p=0.0006. Data plotted as mean ± s.e.m. (E) Quantitation of colocalized VGlut2 & PSD95 synaptic puncta in layer 5 mPFC of wildtype and C3^-/-^ control/CORT mice. Data analyzed by two-way ANOVA, F(1,21)=0.07829, p=0.7824, n=6-7 from 3 mice. Data plotted as mean ± s.e.m. (F) Quantitation of microglial VGlut2 engulfment in layer 5 mPFC of wildtype and C3^-/-^ mice. Data analyzed by two-way ANOVA, F(1,31)=0.6694, p=0.4094, n=8-12 cells from 3-4 mice. Data plotted as mean ± s.e.m.

Finally, we wanted to determine if C3 deletion also leads to the rescue of behavioral deficits. We assessed C3^-/-^ and wildtype mice in two behavioral tasks, the SPT task for anhedonia, and the TOR task for working memory (Figures 7A, B). We found that wildtype mice, but not C3^-/-^ mice, were susceptible to CORT-induced anhedonia in the SPT task (Figure 7C). In addition, we found that wildtype mice but not C3^-/-^ mice were susceptible to CORT-induced impairments in working memory. Specifically, we found that control-treated wildtype mice and CORT/control-treated

**Figure 7.**
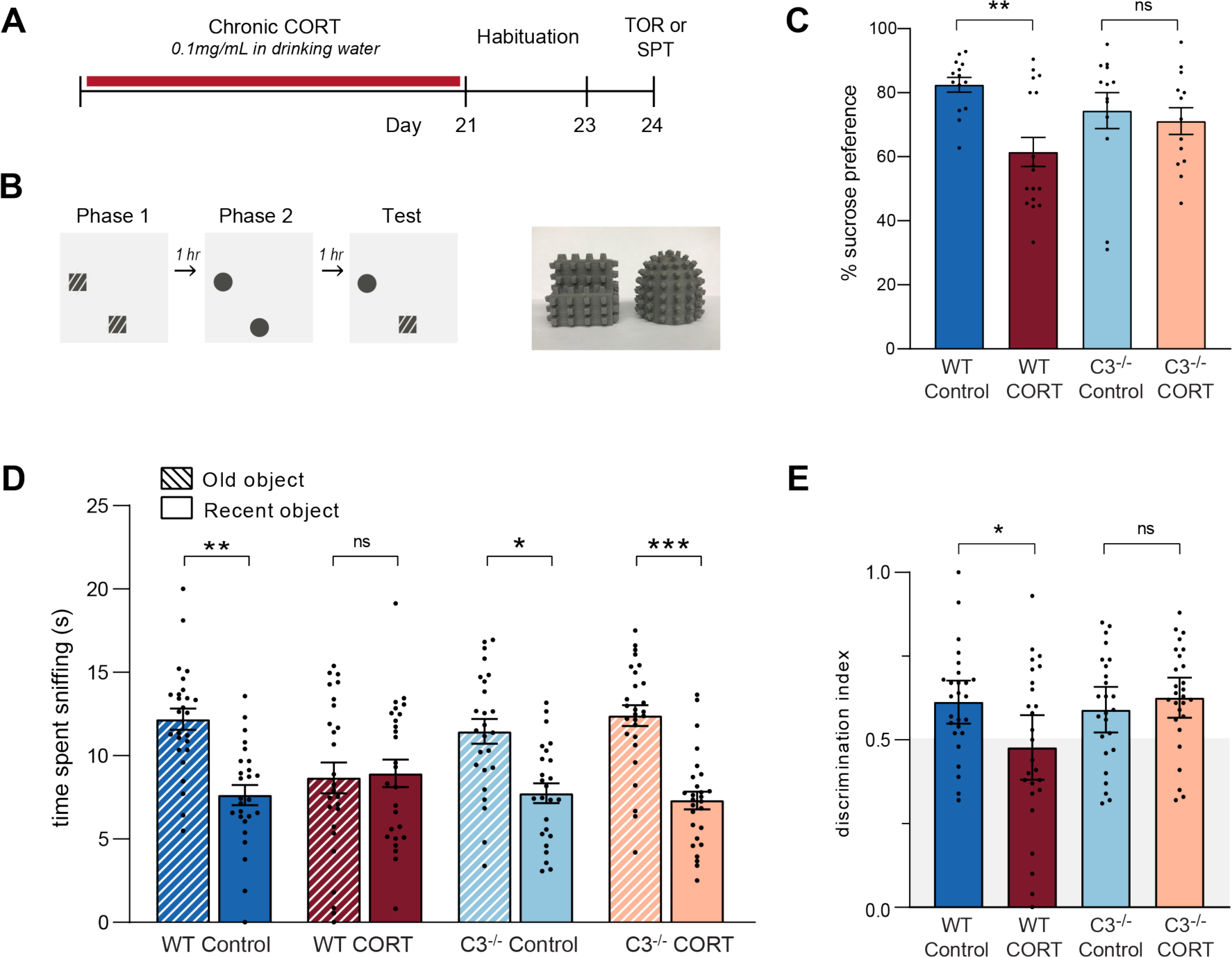
C3^-/-^ mice are protected from stress-induced behavioral deficits. (A) Timeline of CORT administration and behavioral assays (temporal object recognition “TOR” and sucrose preference test “SPT”). (B) Schematic diagram of TOR assay (left), with image of 3D printed objects used for assay (right). (C) Quantitation of sucrose preference of wildtype and C3^-/-^ control/CORT mice. Analyzed by two-way ANOVA, followed by Sidak’s post-hoc test comparing means within genotype, F(1,52)=4.481, p=0.0391 (genotype x stress interaction), n=12-17 per group, wildtype control vs. wildtype CORT **p=0.0017, C3^-/-^ control vs. C3^-/-^ CORT p=0.9298. Data plotted as mean ± s.e.m. (D) Object exploration time during test phase of TOR. Data analyzed by three-way ANOVA, followed by uncorrected Fisher’s LSD test, F(1,100)=5.758, p=0.0183 (object x genotype x stress interaction), n=25-27 per group, adjusted p-values: wildtype control old vs. wildtype control recent **p=0.0018, wildtype CORT old vs. wildtype CORT recent p=0.8376, C3^-/-^ control old vs. C3^-/-^ control recent *p=0.0116, C3^-/-^ CORT old vs. C3^-/-^ CORT recent ***p=0.0004. Data plotted as mean ± s.e.m. (E) Bar graphs showing discrimination index of wildtype and C3^-/-^ control/CORT mice, quantifying relative preference for the old object. Analyzed by two-way ANOVA, followed by Sidak’s post-hoc test comparing means within genotype, F(1,100)=5.784, p=0.0180 (genotype x stress interaction), n=25-27 per group, wildtype control vs. wildtype CORT *p=0.0167, C3^-/-^ control vs. C3^-/-^ CORT p=0.7820.

C3^-/-^ mice were able to remember objects they have recently encountered, as shown by their preferential exploration of the old and more unfamiliar object. In contrast, CORT-treated wildtype mice had impaired working memory and explored both old and recent objects equally (Figures 7D, E). Therefore, our data indicate that C3^-/-^ mice are resilient against stress-induced anhedonia and working memory deficits. Taken together, our findings indicate that stress-induced layer-specific complement activation in the mPFC leads to heterogeneous microglia activation, layer-specific synapse loss and behavioral deficits.

## Discussion

In this study, we show that chronic stress causes spatially-restricted complement activation in the upper cortical layers of the mPFC, resulting in local microglia synapse phagocytosis and synapse loss, as well as anhedonia and working memory deficits. Single cell transcriptomics of the stressed cortex show a dramatic upregulation of the ApoE gene in many microglia, with ApoE^high^ microglia also localized to layer 2/3 of the mPFC. Deletion of the C3 gene blocks the stress-induced layer-specific increases in microglial synapse phagocytosis, synapse loss, and ApoE^high^ microglia population in layer 2/3 of the mPFC. Mice lacking the C3 gene are also resilient against stress-induced anhedonia and working memory deficits. Our findings highlight the role of spatially-restricted microglia activation in the brain in mediating the neurological symptoms of stress-associated diseases such as depression, and reveals the potential role of localized complement and microglia activation in causing region-specific synapse loss in other brain diseases.

The PFC plays an important role in the complex cognitive control of many behaviors, including the regulation of emotion and working memory. During brain development, the PFC is the last region to mature, with an extended developmental period spanning decades in humans^71,72^, making the PFC vulnerable to many environmental and genetic insults. Recent studies indicate that the upper layers of the PFC are preferentially affected in many psychiatric and neurological diseases. Neuropathological studies of brains from schizophrenic patients show a consistent loss of dendritic spines in layer 3 of the dorsolateral PFC^11,12^, while spine density in deeper PFC layers^13^ and other cortical regions^14^ are unaffected. A transcriptomics study of brain cells from autistic patients has also shown that specific sets of genes enriched in neurons and microglia in the upper layers of the PFC are correlated with clinical severity^73^. In Alzheimer’s disease (AD), postmortem brain studies have revealed high Aβ plaque densities in PFC layers 2/3^74–76^, and increased numbers of reactive astrocytes in the same layers^77^. Here, we show that layer 2/3 of the mPFC is selectively vulnerable to stress-induced synapse loss as a result of layer-specific complement and microglia activation. Our findings suggest that the selective vulnerability of the upper layers of the PFC in many brain diseases is based on a propensity for runaway complement activation, although the exact molecular mechanisms and cell types involved remain to be determined.

Many studies have shown that microglia are exquisitely sensitive to the environment. A recent single cell transcriptomic study of microglia has shown that in the somatosensory cortex, microglia adopt layer-specific transcriptional states that are determined by signals from laminarly distributed pyramidal neurons^17^. Therefore, a straightforward mechanism for layer-specific complement activation is the stress-induced increase in the expression of complement components or activators in a particular neuronal or astrocytic layer^78^ in the mPFC. Alternatively, a stress-induced decrease in the expression of complement inhibitors^46,79,80^ within the upper layers can also lead to regional complement activation. In addition, inputs into the mPFC are known to be layer-specific^81^, and a stress-induced signals emanating from specific presynaptic terminals that are laminarly distributed may also trigger layer-specific complement and microglia activation. The above hypothetical mechanisms are not mutually exclusive and may span more than one cell type, and further investigation is required to fully elucidate the mechanisms involved.

In addition to showing that stress induces layer-specific complement and microglia activation, our scRNA-seq data also revealed a stress-induced ApoE^high^ microglia state. We speculate that the ApoE^high^ microglia population phagocytoses synapses in layer 2/3 of the mPFC based on the following lines of evidence: 1) GSEA analysis on the ApoE^high^ microglia state indicate an enrichment of endocytotic genes, suggesting that these microglia perform a phagocytic function; 2) both the ApoE^high^ microglia and the microglia shown immunohistologically to be phagocytosing synapses are localized to layer 2/3, suggesting that they are the same population of microglia, and 3) both the ApoE^high^ microglia and the microglia that phagocytose synapses are ablated by C3 gene deletion, again arguing that they are the same microglia population. However, we acknowledge that these lines of evidence are indirect. We are unable to directly use ApoE as an immunohistological marker for ApoE^high^ microglia because ApoE is also secreted by astrocytes and are avidly phagocytosed by all microglia (data not shown), rendering ApoE staining unreliable as a marker for ApoE^high^ microglia. We were also unable to identify additional genes that simultaneously label the entirety of the ApoE^high^ microglia population and were detectable by high quality antibodies, which precluded the use of alternative markers. Furthermore, the ISH process for detecting ApoE transcripts is moderately destructive on brain slices, and renders them unsuitable for the synapse phagocytosis assay. Therefore, as a result of technical issues, we are unable to directly demonstrate a synapse phagocytosis function for the ApoE^high^ microglia. In addition, it is possible that the ApoE^high^ microglia population may perform other functions, such as cytokine secretion, that are not detected by the DEG and GSEA analyses due to the relatively low sensitivity of the 10x Genomics scRNA-seq platform for low copy number mRNA. Therefore, further work is required to elucidate the biological functions of the stress-induced ApoE^high^ microglia state.

Numerous scRNA-seq studies have shown that microglia adopt disease-associated transcriptomic states. Intriguingly, many of these states include the upregulation of the ApoE gene. Examples of ApoE-associated microglial states include the disease-associated microglia (DAMs) found in AD^21,27^, human AD microglia (HAMs)^82^, the microglial neurodegenerative phenotype (MGnD) found in mouse models of AD/ALS/MS^24^, microglia found in peripheral nerve injury and chronic pain^26^, activated response microglia (ARMs) found in aged mice and an AD mouse model^83^, microglia inflamed in MS (MIMS)^25^, and a microglia state associated with injury^19^. The upregulation of microglial ApoE in a wide variety of diseases suggests that ApoE upregulation is part of a generic microglial response to tissue damage. Interestingly, early studies in stress biology describe the stress response as a “general adaptation syndrome” that occurs in diverse diseases^84,85^. In this study, we show that the stress hormone corticosterone induces an ApoE^high^ microglia state, suggesting that the ApoE component of multiple disease-associated microglial states may be a result of stress hormone release in those diseases.

Stress is also known to aggravate or trigger the occurrence of other neurological and psychiatric diseases. For example, stress is known to trigger episodes of psychosis in schizophrenic patients^86,87^ and attacks in multiple sclerosis patients^88^, as well as accelerate the progression of AD^89–91^. All of these diseases are associated with aberrant complement activation in the brain^25,92–99^. Our findings suggest that stress may aggravate these diseases through additive complement and microglia activation. We also show that only VGlut2 synapses in layer 2/3 mPFC are phagocytosed by microglia, while VGlut1 and VGAT synapses are spared, indicating that even within a region of complement activation, specific microcircuits may be selectively impaired. These findings may be relevant when assessing the neural circuits involved in various complement-associated brain diseases.

Neuroimaging studies have revealed disease-specific and reproducible patterns of gray matter volume (GMV) reductions in many brain diseases, including depression^100–102^, schizophrenia^103^, and other major psychiatric diseases^104,105^. A more recent study has showed that substantial heterogeneity also exists in the individual pattern of GMV changes in patients diagnosed with the same psychiatric disorder, and suggests that individual variation in disease symptoms may be a result of this heterogeneity^104^. Other studies have also shown that elevated salivary, hair and serum cortisol levels are all correlated with lower brain volumes^38,39^, and that stressful life events leave a replicable pattern of brain volume reductions that is predictive of current and future anxiety^106^. While GMV is a complex measure that encompasses glia, vasculature and neuronal dendritic and synapse volumes, longitudinal studies show a lifetime trajectory of GMV loss^107,108^ that temporally correlates with neuropathological studies of dendritic spine loss due to increased synapse pruning during adolescence and early adulthood^109,110^, suggesting that synapse loss is a major contributor to GMV reductions. Given the large variation in complement activity found in the human population^111^, genetic polymorphisms in complement components and regulators may account for the individual variation in GMV reductions observed in psychiatric patients. Our study suggests that lifetime adversity can leave spatially heterogeneous patterns of brain volume reductions through heterogeneous complement and microglia activation, and that regional regulation of these processes play an important role in the heterogeneity of neuropsychiatric diseases.

## Acknowledgements

This work is supported by the National Institute of Neurological Disorders and Stroke (NINDS) grant (R01NS112389) to G.M.S., the National Institute on Deafness and other Communication Disorders (NIDCD) grants (R01DC018797, R01DC019371) to J.H.K, the National Institute of Mental Health (NIMH) grant (R01MH053851) to D.A.M., and the NIMH grant (R21MH113899) to F.R.C.. Imaging was performed at UTHSA Optical Imaging Facility, which is supported by a grant from the NCI (P30CA54174). FACS was performed at the Flow Cytometry Shared Resource at UTHSA, which is supported by an NCI grant (P30CA054174) to the Mays Cancer Center, a Cancer Prevention and Research Institute of Texas (CPRIT) grant (RP210126), a National Institutes of Health (NIH) Shared Instrument grant (S10OD030432), and support from the Office of the Vice President for Research at UTHSA. RNA sequencing was performed at the UTHSA Genome Sequencing Facility, which is supported by an NCI grant (P30CA054174), an NIH Shared Instrument grant (S10OD030311), and a CPRIT core facility award (RP220662).

## Author contributions

G.M.S. and B.M.S. designed the study. B.M.S. carried out most of the anatomical and behavioral experiments. H.T., H.C., and J.G. assisted with anatomical and behavioral experiments. M.W. and J.H.K. assisted with neuronal functional experiments. J.B.L. assisted with scRNA-seq analysis. D.A.M. and F.R.C. assisted with the design of the stress protocols. G.M.S. supervised the project. G.M.S. and B.M.S. wrote the manuscript.

## Declaration of interests

The authors declare no competing interests.

## Data availability

Single cell RNA sequencing data will be deposited at the Gene Expression Omnibus (GEO) and be publicly available upon publication. All additional data and code will be provided upon request.

---

**Supplemental Figure 1.**
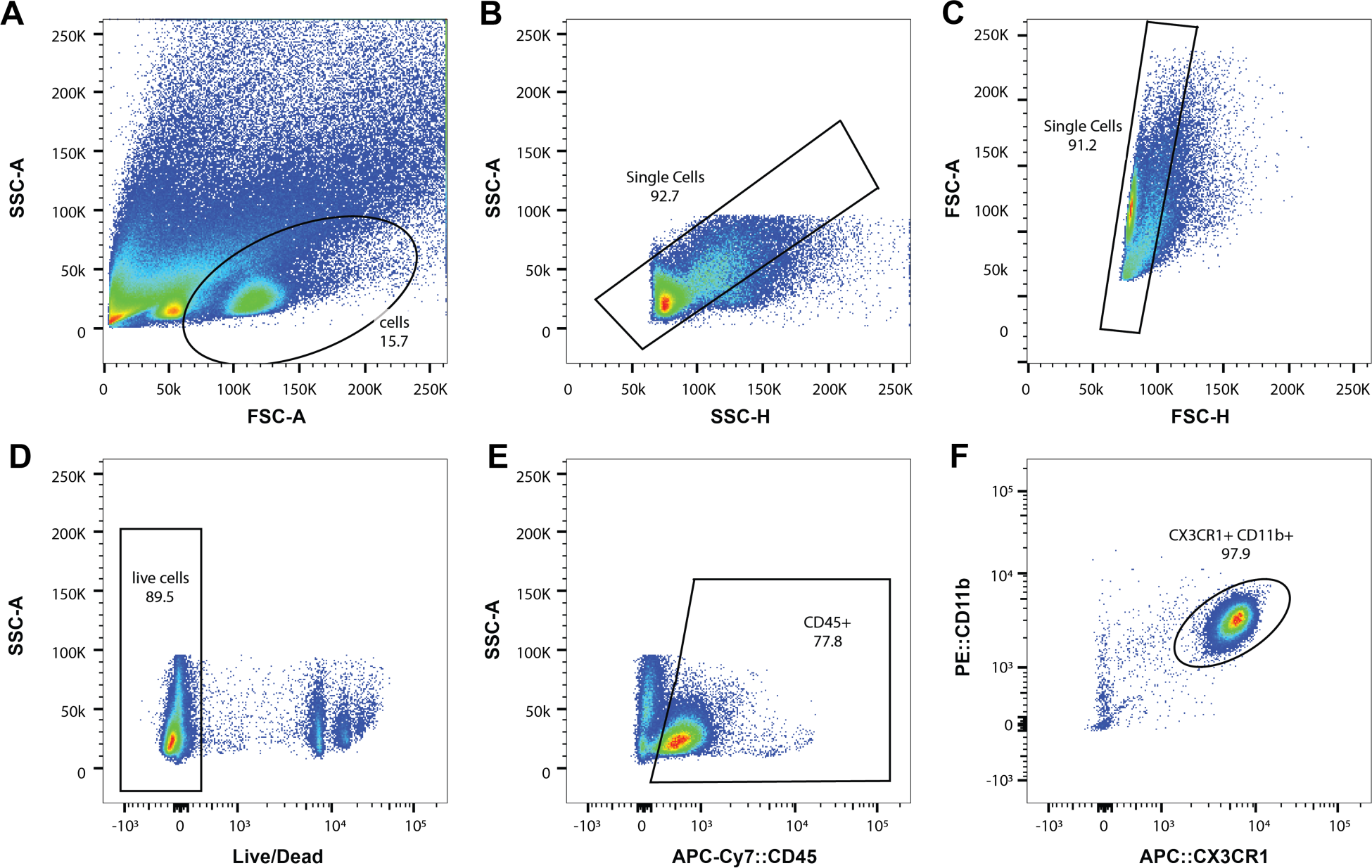
FACS gating strategy for microglia isolation before scRNA sequencing. (A) Representative FACS plot of input cell suspension, and separation of cells from debris based on forward and side scatter. (B) Representative FACS plot for isolation of single cells based on side scatter. (C) Representative FACS plot for isolation of single cells based on forward scatter. (D) Representative FACS plot for isolation of live cells based on low DAPI staining. (E) Representative FACS plot for isolation of CD45^high^ microglia cells. (F) Representative FACS plot for isolation of Cx3cr1^high^ and Cd11b^high^ microglia cells.

## Methods

### Animal care

All animal work was approved by the UT Health Institutional Animal Care and Use Committee (IACUC). Colonies were maintained in climate-controlled rooms within the AAALAC-accredited UT Health animal facility. Mice were group-housed (3-4 per cage) under a 14-hour light/10-hour dark cycle with ad libitum access to food and water. Clear red-tinted tunnels (Bioserv, K3322) were placed in all cages for enrichment and transport purposes. To reduce distress during transfer, mice were habituated to tunnel handling with daily 3-5 min tube rides for 1 week prior to behavioral studies. Behavioral assays were conducted either during the last 4 hours of the light cycle (temporal object recognition) or throughout the dark cycle (sucrose preference test).

### Chronic corticosterone (CORT) stress protocol

Mice received either corticosterone-infused (CORT) drinking water (Sigma 27840, corticosterone dissolved at 10 mg/ml in 100% ethanol, then mixed with facility-provided drinking water to a final concentration of 0.1 mg/ml corticosterone and 1% ethanol) or control drinking water (1% ethanol) for 21 days, as described^54^. Both CORT and control water were prepared in standard bottles using autoclaved drinking water from the animal facility and replaced weekly.

### Chronic restraint stress (CRS) protocol

Mice were subjected to 4 hours of restraint stress daily for 21 days, as described^54^. Restraint tubes were made from 50 mL canonical tubes (Falcon, #2098) with drilled holes for air circulation. Restraint tubes were placed inside the home cage. Control mice were handled twice daily, 4 hours apart, for 5 minutes each. Tubes were cleaned with 70% ethanol between uses.

### Chronic unpredictable stress (CUS) protocol

We used a protocol that had been shown in multiple studies to cause stress-induced circuitry changes in mice^112,113^. Mice were subjected to two different stressors per day, once in the morning and once in the afternoon, for 10 days. Stressors included: 15 min tail pinch, 2 hr tube restraint, 30 min elevated platform, 10 min inescapable footshock (0.7mA, 2 s, 10 total shocks), 24 hr wet bedding with 45° cage tilt, and 24 hr light exposure. Control mice were handled twice daily. CUS mice were singly housed for the duration of the stress protocol, while control mice were group housed.

### Immunohistochemistry and microscopy

Mice were deeply anesthetized with Avertin (0.5 mg/g) for transcardial perfusion with ice-cold PBS and 4% paraformaldehyde (PFA). Brains were rapidly extracted, post-fixed for 2 hrs at 4°C with gentle rocking, then cryoprotected in 30% sucrose in PBS for 48-72 hours. Brains were sliced on a sliding-freezing microtome (Leica) into 20 μm sections (for synapse and complement staining) or thick 40 μm sections (for microglia staining). Wash, block and antibody solutions were made from PBS (pH 7.4) with either 0.15% TX-100 (0.15% PBT) or 0.3% PBT, depending on slice thickness. Free-floating sections were washed 2x in PBT (10 min/wash), post-fixed in 4% PFA (15min) and briefly rinsed in PBS (5 min) while rocking at room temperature (RT). Sections were blocked in 10% normal donkey/goat serum (Jackson Immunoresearch) at RT for 2 hr, then incubated in primary antibodies in 1% blocking buffer overnight at 4°C with gentle rocking. Once equilibrated to RT, sections were washed 4x (10 min/wash), then incubated in secondary for 2 hr. Sections were washed 2x in PBT (5 min/wash), 3x in PBS (10 min/wash) then floated onto glass slides. Slides were treated with TrueBlack Lipofuscin Autofluorescence Quencher (Biotium, 23007), rinsed 3x in PBS (3min/wash), then cover-slipped with Fluoromount-G (SouthernBiotech, 0100-01). Slides were stored in 4° and imaged within 1 week.

The following primary antibodies were used for staining: goat anti-C3 (1:1000, MP Biomedicals, cat#: 855730), rabbit anti-Iba1 (1:500, Fuji Film, cat#: 019-19741), rat anti-CD68 (1:500, Bio-Rad, cat#: MCA1957), guinea pig anti-VGlut2 (1:1000, Synaptic Systems, cat#: 135 404), rabbit anti-VGlut1 (1:1000, Synaptic Systems, cat#: 135 302), mouse anti-PSD95 K28/43, IgG2a (1:500, Neuromab, cat#: 75-028), guinea pig anti-VGAT (1:500, Synaptic Systems, cat#: 131 004), mouse anti-gephyrin, IgG1 (1:500, Synaptic systems, cat#: 147 021), rabbit anti-cFOS (1:500, Synaptic systems, cat#: 226-008).

The following secondary antibodies were used for staining: donkey Anti-Goat IgG A488 (1:500, Jackson, cat#: 705-545-147), donkey Anti-Rabbit IgG A488 (1:500, Jackson, cat#: 711-545-152), cy3-AffiniPure Donkey Anti-Rat IgG H&L (1:500, Jackson, cat#: 712-165-153), goat Anti-Guinea Pig IgG Polyclonal CF-488A (1:500, Biotium, cat#: 20017-1), cy5-AffiniPure Donkey Anti-Guinea Pig IgG H&L (1:500, Jackson, cat#: 706-175-148), goat Anti-Rabbit IgG H&L Cy5 (1:500, Abcam, cat#: ab6564), goat anti-Mouse IgG2a A555 (1:500, Thermo Fisher Scientific, cat#: A-21137), goat anti Guinea Pig IgG H&L A647 (1:500, Thermo Fisher Scientific, cat#: A-21450), goat anti-Mouse IgG1 A488 (1:500, Thermo Fisher Scientific, cat#: A-21121).

Sections were imaged using a Zeiss AxioObserver inverted microscope equipped with a Zeiss Apotome.2 module. Images were acquired using a Zeiss 63x/1.4 oil Plan-Apochromat objective (synapse density, microglial activation and engulfment, high-resolution C3), or Zeiss 20x Plan-Neofluar objective (low-magnification C3 staining). Identical imaging and analysis parameters were used for each experimental group.

### Image analysis for C3 deposition and synapse densities

ImageJ (NIH) was used to quantify synapse densities and C3 deposition. Individual channels were first Z-projected with summation over a depth of 2 µm, followed by background subtraction (8 px rolling ball radius) and gaussian filter (sigma = 0.8). The threshold was empirically determined for each channel and applied consistently to all groups. Watershed was applied to binarized image. For synapse density, colocalized puncta were revealed by multiplying the individual pre & post-synaptic channels using the image calculator tool. The Analyze Particles tool was used to quantify the number of colocalized synaptic puncta and individual C3 deposits.

### Image analysis for microglial lysosome content and synapse phagocytosis

Reconstructions of microglia were performed using high-resolution confocal-quality images of Iba1-positive cells using Imaris Software (version 9.7.1, Bitplane, Concord MA), as previously described^45,46^. Using the image processing tool, background subtraction was first performed (sigma = 1 for VGlut2, CD68; sigma = 60 for Iba1), followed by the application of a gaussian filter (sigma = 0.144). All surfaces were rendered with 0.1 µm smoothing. Iba1-positive cells were reconstructed first, and disconnected processes were unified with the cell body to create a single surface representing one cell. To render CD68, a mask was then applied to isolate the CD68 signal only within the reconstructed microglia. Another mask was then applied to isolate the VGlut2 signal only within the reconstructed CD68-positive lysosomes within microglia. Volumes were recorded for each reconstructed surface. For microglial lysosome content, the percent of CD68 occupancy within microglia was calculated using the formula: *(volume of CD68/volume of Iba1+ cell)*100*. For microglial engulfment of VGlut2, the percent of VGlut2 within microglial lysosomes was calculated using the formula: *(volume of VGlut2 within lysosome/volume of Iba1+ cell)*100*.

### Quantification of c-fos positive cells

Mice were sacrificed exactly 1 hour after completion of the TOR test phase. Brain tissue was collected and processed as described above with slight modifications. In short, 40 µm-thick brain sections were fixed in ice-cold 100% methanol for 5 minutes, washed in 0.3% PBT for 15 minutes, then blocked in 1% BSA 5% NDS in 0.3% PBT for 2 hours at RT. Sections were incubated in primary antibody in blocking buffer overnight (rabbit anti-c-fos, 1:1000, Synaptic Systems 226-008), washed 5x with PBT, then incubated in secondary antibody (donkey anti-rabbit A488, Jackson, catalog #: 711-545-152) for 2 hr at RT. Sections were imaged on a Zeiss AxioObserver inverted microscope equipped with Apotome.2 module using a Zeiss 20x Plan-Neofluar objective. Analysis of c-fos positive cells was automated using an ImageJ macro by applying the following sequence on 200 µm x 200 µm ROIs: gaussian filter (sigma = 2), threshold (1000,65535), convert to binary mask, watershed, analyze particles.

### Microglia single cell dissociation and FACS

Microglia single cell dissociation was performed using the cold mechanical dissociation procedure which minimizes ex vivo activation of microglia as previously described^19,65^. Brains were collected from control and CORT mice after transcardiac perfusion of ice-cold HBSS, and the cerebral cortices were dissected out and minced after removal of white matter. Pieces of cortices were then Dounce homogenized for 20 strokes with a loose rotating pestle, and then strained through a 70 µm strainer. Cells were washed and pelleted in HBSS, and then resuspended in 40% Percoll. Cells were spun at 500 g in the cold for 1 hr, and the pelleted cells were washed in HBSS and resuspended in FACS buffer for staining. Cell suspensions were incubated with Fc block reagent (1:50 dilution) for 15 min on ice, and then the following antibodies were added from a master mix: DAPI (1:10,000), Cd11b-PE (1:50), CD45-APC-Cy7 (1:100), Cx3cr1-APC (1:250). Cell suspensions were further incubated for 30 min on ice, and then washed twice and resuspended in FACS buffer. Cells were then immediately transported to the Flow Cytometry Core Facility on ice for sorting. Sorting was performed with the following gating strategy (Supplementary Figure 1): whole cells versus debris (forward scatter (FSC-A) vs side scatter (SSC-A)), singlets vs aggregates (SSC-H vs SSC-A, and FSC-H vs FSC-A), live vs dead (DAPI vs SSC-A), Cd45^high^ (APC-Cy7-CD45 vs SSC-A), Cx3cr1^high^ and Cd11b^high^ (APC-CX3CR1 vs PE-CD11b).

### Single cell library generation and sequencing

After sorting in the cold, cells were put on ice and immediately transported to the Genome Sequencing Core Facility for 10X Genomics single-cell capture and sequencing with the Chromium 3’ library v3.1 kit, following manufacturer’s instructions. Approximately 10,000 cells were targeted per sample for single cell sequencing. The single cell libraries were then sequenced on an Illumina NovaSeq6000 sequencer, targeting at least 500M reads per sample, for an average sequencing depth of 50,000 reads per cells. Cell Ranger v7.0.1 (10X Genomics) was used for sample demultiplexing, barcode processing, unique molecular identifiers filtering, gene counting, and mapping to the mouse mm10-2020-A (10X Genomics) reference genome.

### scRNA-seq clustering analysis

Single cell clustering analysis was performed with the R package Seurat v4.3.0^114^. For initial quality filtering, genes expressed in more than 5 cells were kept. Cells with greater than 5% mitochondrial gene expression or less than 2% ribosomal gene expression were then removed. Lastly, cells expressing more than 2000 genes and 5000 UMIs were kept. The SCTransform v2 algorithm^115^ from Seurat was then used to normalize the counts in each sample. Cells were then filtered for doublets using the R package DoubletFinder^116^, based on an expected doublet rate of 7.6% for 10,000 cells per channel, as specified in the manufacturer’s instructions. After filtering, a total of 35,292 microglia were retained for downstream analysis.

Cells from all samples were integrated with Canonical Correlation Analysis (CCA) with Seurat’s *SelectIntegrationFeatures*, *PrepSCTIntegration*, *FindIntegrationAnchors*, and *IntegrateData* functions. Principal Components Analysis (PCA) was then performed for the integrated dataset, and the top 20 principal components (PCs) were chosen for clustering based on an elbowplot. Clustering was performed using the *FindNeighbors* function with 20 PCs, and the *FindClusters* function at a resolution of 0.08. Uniform Approximation and Projection (UMAP) dimensionality reduction was performed to visualize the cell clusters with the *RunUMAP* and *DimPlot* functions. Marker genes for clusters were identified using the *FindAllMarkers* function with the Wilcoxon rank-sum test. The top expressing genes in each cell cluster was plotted as a heat map using the *DoHeatMap* function. Comparison of differentially expressed genes (DEGs) between the homeostatic and ApoE^high^ clusters were performed with the *FindMarkers* function with the parameter max.cells.per.ident = 3000, and the results were plotted in a volcano plot using the EnhancedVolcano R package^117^.

### Pseudotime trajectory analysis

The R package Monocle3^118^ was used to construct a pseudotime trajectory in the clustered dataset and to identify genes that were differentially expressed along the pseudotime trajectory. The *learn_graph* function of Monocle3 was used to fit a principal graph in the combined dataset, and the *order_cells* function was used to order the cells into trajectory along the principal graph, with graph nodes in the homeostatic cluster specified as the root of the trajectory. Genes that were differentially expressed along the pseudotime trajectory were determined using the *graph_test* function.

### Gene set enrichment analysis (GSEA)

The R package clusterProfiler^119^ was used to perform GSEA. The list of DEGs from the pseudotime trajectory analysis (log2FC > 0.25 and adjusted p value < 0.05, resulting in a total of 93 genes) was used as input to the *enrichGO* function of clusterProfiler with the following parameters: keyType = “SYMBOL”, OrgDb = org.Mm.eg.db, ont = “BP”, pvalueCutoff = 0.05, pAdjustMethod = “BH”. The top 10 enriched biological process (BP) gene ontology (GO) terms were ranked in order of increasing adjusted p value, and plotted as a dot plot.

### RNAscope ISH

RNAscope™ Multiplex Fluorescent V2 assay (ACDBio, 323136) was carried out according to manufacturer’s instructions, with some modifications. Briefly, brains from PBS-perfused mice were rapidly dissected and snap frozen in liquid nitrogen. Brains were sliced using a cryostat into 16um sections and adhered onto super-frost plus slides (Electron Microscopy Sciences, 71869-20). Slides were kept on dry ice during slicing and then promptly stored at -80°C in a sealed box until use. Immediately upon removal from the freezer, slides were plunged into fresh ice-cold 4% PFA for 15 min, rinsed 2x with PBS, then dehydrated in serial dilutions of ethanol (50%, 75%, 100%, 100%). Slides were then air-dried for 5 min, and a PAP pen was used to create a hydrophobic barrier around the tissue. For all subsequent steps, slides were moved to the HybEZ Humidity Control Tray with wet humidifying paper for treatment/incubation, and washed in coplin jars with gentle rocking (2 min per wash). At room temperature (RT), sections were treated with hydrogen peroxide for 10 min, washed 3x in distilled water, treated with Protease IV for 20 min, then washed 3x with PBS. Probes (Mm-CX3CR1-C2, cat no. 314221-C2 and ApoE-C3, cat no. 313271-C3) were hybridized to tissue at 37°C for 2 hr, rinsed 3x in wash buffer, and stored in 5x saline sodium citrate (5x SSC) buffer overnight at RT. Slides were rinsed 1x with wash buffer before experiment continuation. Subsequent amplification & HRP steps were performed according to manufacturer’s instructions. HRP-C2 and HRP-C3 were labelled with Opal 520 (1:750, Akoya FP1487001KT) and Opal 570 (1:1500, Akoya FP1488001KT), respectively. Slides were counterstained with DAPI, cover slipped with Prolong Gold (Cell Signaling Technology #9071), stored in 4°C and imaged within 1 week.

Images were acquired with a Zeiss AxioObserver equipped with an Apotome.2 module using a Zeiss 40x Plan-Neuofluar objective. Analysis of ApoE expression was limited to CX3CR1+ DAPI nuclei. ApoE expression within microglia was evaluated using the ACDBio semi-quantitative scoring system: “0” represents no staining or less than one dot per ten cells, “1” represents one to three dots per cell, “2” represents four to nine dots per cell and none or very few dot clusters, “3” represents 10 to 15 dots per cell and/or less than 10% of dots in clusters, and “4” represents more than 15 dots per cell and/or more than 10% of dots in clusters. Bin counts were aggregated within each experimental group (N=3-4 mice per group, 1-2 images per mouse) to determine the bin distribution for each condition.

### Sucrose preference test (SPT) task

Sucrose preference test was conducted using a 1% sucrose solution (Sigma S9378-1KG). Bottles were made from 50 mL conical “iTubes” with low-aldehyde odor (Stellar Scientific, T50-600) custom fitted with 2.5” sipper tubes (Fisher, NC1636707) and rubber stoppers (VWR, 76293-684) to minimize leakage. Sucrose and control water were prepared using autoclaved drinking water from the animal facility. Sucrose water and control water were sterile filtered before aliquoting into conical tubes. Mice were simultaneously habituated to sucrose and control water bottles for 3 consecutive days prior to testing. Bottles were switched in positions once daily to avoid side preference. On the test day, mice were separated into individual housing and deprived of water for the last 4 hours of the light cycle. Sucrose and water bottles were weighed at the beginning and end of the dark cycle. Percent sucrose preference was calculated with the following formula: *((sucrose consumption - water consumption) / total consumption) x 100*

### Temporal Object Recognition (TOR) task

TOR was conducted in an 80 x 80 x 30 cm white acrylic open field arena, with white opaque wall inserts separating the arena into (4) 40 x 40 x 30 cm height square chambers (Conduct Science). Objects were 3D printed from natural PLA filament (3D printer files provided by Maithe Arruda-Carvalho^120^). Two types of objects were printed: steps and domes. To minimize intrinsic preference of mice for either object, both objects were made of the same material, and were designed to be equal in surface area. Objects were also tested in pilot studies to verify that mice had no intrinsic preference for either object, and counterbalanced between phase 1 and phase 2 as described below. Objects were centered midway along the inside of the external chamber walls, equidistant from the corners. To prevent movement, objects were affixed to the chamber floor using a small piece of BluTack adhesive putty (a fresh piece was used every round for every mouse). Chambers and objects were cleaned with 15% ethanol between each mouse and phase to minimize odor carryover. All equipment was cleaned with 70% ethanol at the end of the test session. Mice were habituated to each quadrant (chamber) of the arena, 5 min per chamber, for two consecutive days prior to testing. The TOR procedure consisted of three 5-minute phases (phase 1, phase 2 and test) with 1 hr inter-phase intervals. Mice were returned to home cage between phases. During phase 1, mice were placed in the arena with a pair of identical objects (either 2 steps or 2 domes). After a 1 hr delay, mice were reintroduced to the same chamber containing a different pair of identical objects for phase 2. During the test phase, mice were reintroduced to the same chamber containing one object from phase 1 (old object) and one object from phase 2 (recent object). Sessions were recorded using an infrared camera (OEM Cameras, DFK 33UX290) with a 3-8mm focal length lens (OEM Cameras, T3Z0312CS-MPIR) fixed to the ceiling directly above the arena. Videos were analyzed using Noldus Ethovision tracking software (Noldus Ethovision XT, Version 11.5). Behavioral analysis during the test phase was restricted to the first 20 s of total interaction time with objects, as previously described^120^. The object discrimination index was calculated using the following formula: *old object interaction time / total object interaction time*.

## Notes

### Competing Interest Statement

The authors have declared no competing interest.

### Summary of Updates

Manuscript text has been revised to reflect reviewer feedback.

